# Comparing orthology methods and their performance by recapitulating patterns of eukaryotic genome evolution

**DOI:** 10.1101/2020.05.13.092791

**Authors:** Eva S. Deutekom, Berend Snel, Teunis J.P. van Dam

## Abstract

Insights into the evolution of ancestral complexes and pathways are generally achieved through careful and time-intensive manual analysis often using phylogenetic profiles of the constituent proteins. This manual analysis limits the possibility of including more protein-complex components, repeating the analyses for updated genome sets, or expanding the analyses to larger scales. Automated orthology inference should allow such large scale analyses, but substantial differences between orthologous groups generated by different approaches are observed.

We evaluate orthology methods for their ability to recapitulate a number of observations that have been made with regards to genome evolution in eukaryotes. Specifically, we investigate phylogenetic profile similarity (co-occurrence of complexes), the Last Eukaryotic Common Ancestor’s gene content, pervasiveness of gene loss, and the overlap with manually determined orthologous groups. Moreover, we compare the inferred orthologies to each other.

We find that most orthology methods reconstruct a large Last Eukaryotic Common Ancestor, with substantial gene loss, and can predict interacting proteins reasonably well when applying phylogenetic co-occurrence. At the same time derived orthologous groups show imperfect overlap with manually curated orthologous groups. There is no strong indication of which orthology method performs better than another on individual or all of these aspects. Counterintuitively, despite the orthology methods behaving similarly regarding large scale evaluation, the obtained orthologous groups differ vastly from one another.

**Availability and implementation:** The data and code underlying this article are available in github and/or upon reasonable request to the corresponding author: https://github.com/ESDeutekom/ComparingOrthologies.

**Summary:**

- We compared multiple orthology inference methods by looking at how well they perform in recapitulating multiple observations made in eukaryotic genome evolution.
- Co-occurrence of proteins is predicted fairly well by most methods and all show similar behaviour when looking at loss numbers and dynamics.
- All the methods show imperfect overlap when compared to manually curated orthologous groups and when compared to orthologous groups of the other methods.
- Differences are compared between methods by looking at how the inferred orthologies represent a high-quality set of manually curated orthologous groups.
- We conclude that all methods behave similar when describing general patterns in eukaryotic genome evolution. However, there are large differences within the orthologies themselves, arising from how a method can differentiate between distant homology, recent duplications, or classifying orthologous groups.

## Introduction

Gaining insight into the evolution of eukaryotic pathways and protein complexes is often obtained by careful and intensive manual efforts of inferring orthologous groups (OGs), often using phylogenetic profiles [1–4]. The reconstructions of the evolution of these pathways is changing our view of eukaryotic genome evolution. With the increase of genomic data, specifically of divergent eukaryotes, the pervasiveness of gene loss became a clear pattern for genetic variation [5–7] that has been observed in many eukaryotic functional pathways, protein complexes and metabolic pathways [1–4,8,9]. From these manual reconstructions it is becoming more apparent that the loss of parts of these complexes or pathways between the Last Eukaryotic Common Ancestor (LECA) and extant species is mostly non-random, as whole (sub)complexes are often co-lost or, at least, co-absent. A final theme in these studies is that more and more processes and pathways are inferred to have likely been present in LECA and thus we are faced with an ever expanding LECA [10].

In addition to the insights offered for the study of specific pathways, large scale orthologies should provide benefits that overcome the limits in case studies. Manually curated OGs and phylogenetic profiles are laborious and don’t scale well when new or updated genomes become available. The development of accurate computational methods to automatically infer orthologies to the same degree as manual analysis is challenging since the orthology inference algorithms need to take into account complex histories of genes, such as duplications, losses and gains of genes and/or their domains and horizontal gene transfer [11–14]. Readily available curated orthology databases are often very useful for more specific evolutionary questions, e.g. TreeFam database containing mainly animal gene families [15], or OrthoDB containing mainly vertebrate, arthropod, fungi, plant and bacterial gene families [16]. For other evolutionary questions these databases are not always practical, as the species covered in these databases might not be suitable for a particular evolutionary question, e.g. due to (limited) taxonomic range or over/under-sampling of species.

In an effort to progress this field, a large group of researchers have come together to work on a collaborative quest for orthologs (QfO) and have derived a suite of benchmarking tools to determine the performance of orthology methods, both old and new, in a systematic manner [17]. The QfO Benchmarking suite is a very powerful way to objectively evaluate orthology methods in a generic way and has thereby paved the way for advancement in the field of orthology. QFO focusses on a single metric, namely how well an orthology method can recapitulate OGs in a gold standard fixed set of reference proteomes. The benchmark results of orthology methods thus depend on this reference set.

Complementary to the QFO, we here investigate different orthology inference methods and how well they can recapitulate a number of observations made regarding genome evolution in a large diverse set of eukaryotic genomes. We specifically investigate loss patterns and numbers, co-occurrence and LECA gene content. We expand our analysis by investigating qualitative differences between the obtained orthologies and their inferred OGs, and how they capture high quality manually curated OGs.

We find that most orthology methods reconstruct a large LECA, with substantial gene loss, and can predict interacting proteins reasonably well when applying phylogenetic co-occurrence. However, derived OGs show imperfect overlap with manually curated OGs. There is no strong indication of which orthology method performs better than another on either or all of these aspects. Counterintuitively, despite the orthology methods behaving similarly regarding large scale evaluation, the obtained OGs differ vastly from one another.

## Methods and Materials

### Inferring (LECA) orthologs in a large scale dataset

To investigate different automated orthology methods we inferred orthologous groups (OGs) with 167 proteomes (2865661 sequences) from a diverse set of eukaryotes [18]. Since we have a large set of proteomes, the orthology inference methods used in this study were chosen based on reasonable computational time, ease of parallelizing the process on multiple CPU’s, ease of projection on our set of proteomes, and development activity. Also, the methods must be able to infer OGs of genes: multiple orthology inference methods we found have pairwise species comparisons, and not the multispecies comparisons that give OGs. Table 1 lists the orthology inference methods and databases chosen for this study and a brief description of each. There are other orthology inference methods that were considered [19–22], but due to various reasons were not used in this study. They are listed in Supplementary Table 1 with a short description.

**Table 1.** Orthology inference methods and databases used in this study.

| Tool/Dataset | Prediction type | Description and notes |
| --- | --- | --- |
| Ancestral Panther | Database | Ancestral genomes dataset contains reconstructed ancestral genomes based on gene family trees from the PANTHER database, from which HMM profiles were built. |
| Broccoli | De novo prediction | K-mer pre-clustering to simplify proteomes, followed by a similarity search (DIAMOND) and phylogenetic analysis (FastTree2). Orthologous groups are inferred using a machine learning algorithm, LPA. Extremely fast when run on a large dataset. |
| EggNOG (DIAMOND and hmmer) | Database | Manually curated sequence sets ran with (1) seed ortholog assignments (DIAMOND) and (2) HMM profile searches (hmmer). |
| Orthofinder (DIAMOND and BLAST) | De novo prediction | Uses both (1) DIAMOND or (2) BLAST as an aligner. Has a sequence length and phylogenetic distance normalized bit-score cut-off between pairs of genes, which function as edge weights in the orthogroup graph. Clustering of genes is done with the MCL method. |
| SonicParanoid | De novo prediction | Uses MMseqs2 as an aligner. The algorithm of InParanoid is used as backbone, with changes to the core algorithm that reduce the execution time and increase the usability of the tool. Relies on cumulative alignment score of groups and avoids using thresholds based on confidence score between pairs of genes. Clustering of genes is done with the MCL method. |
| SwiftOrtho | De novo prediction | Taking the same approach as OrthoMCL for normalized bit-score cut-off between pairs of genes, which function as the edge weights in the graph. Clustering of genes is done with the MCL method. SwiftOrtho is optimized for speed and memory usage when applied to large-scale data. |

The eukaryotic eggNOG [23] hmm profile database (version 4.5.1) was used to annotate our eukarya dataset with hmmsearch (version 3.2.1 | June 2018 in HMMER 3.1b2 package February 2015) and a hit cut-off e-value of 10^-3^. Additionally, the eggNOG protein database (version 4.5.1) was used to annotate the eukarya dataset with DIAMOND and a hit cut-off e-value of 10^-3^. For both strategies we used the emapper annotation tool (version emapper-1.0.3) provided by eggNOG. We wanted to see if there exist any (large) differences between these strategies.

We applied Orthofinder (version 2.1.2) [24] with default values using both all-vs-all BLAST (hit cut-off e-value 10^-3^) and DIAMOND (hit cut-off e-value 10^-3^) as sequence aligners. We chose DIAMOND as a high-throughput aligner that is multiple orders of magnitude faster than BLAST [25]. Also, here we wanted to see if there exist any (large) differences between the alignment strategies.

We additionally used Broccoli (version 1.0) [26] with default parameter settings. SonicParanoid (version 1.3.0) [27] was used with default parameter settings in the sensitive mode (MMseqs2 sensitivity parameter is set to s = 6.0 by SonicParanoid), which is suggested for more distantly related species such as our diverse set of eukaryotes. Since we were not able to get SonicParanoid running on our device, we obtained the results through personal communication with the authors. SwiftOrtho [28] was run using all-vs-all BLAST (hit cut-off e-value 10^-3^) and the default parameter settings.

In order to investigate loss patterns for each orthology and to manage the amount of orthogroups in further analysis steps, following the orthogroup inference by each method we used the Dollo parsimony approach [29–31] with an additional strict inclusion criteria [18] to infer orthogroups that were likely present in LECA. Briefly, the Dollo parsimony method assumes genes can be gained only once and losses are minimized. To be called a LECA OG, the genes belonging to the OG must be in at least three eukaryotic supergroups distributed over opimoda and diphoda species.

Additionally, we used Panther ancestral genes from the Ancestral Genomes Resource database [32] that has ancestral genes up to the Last Universal Common Ancestor. Gene trees and corresponding multiple sequence alignments were obtained through personal communication with the authors. We acquired eukaryotic ancestral genes by traversing the gene trees with an inhouse script using the ete3 package and obtaining genes in the eukaryotic ancestral nodes, the leaves of these ancestral nodes we define as LECA genes. Additionally, we followed the same criteria for the other orthologies above to select LECA genes and require inclusion of at least three eukaryotic supergroups distributed over both opimoda and diphoda species to avoid including possible issues with in/outparalogy resulting from erroneous tree inference. We trimmed the provided multiple sequence alignments with an inhouse script using the biopython package to remove empty columns that were left after obtaining LECA genes. Next, we made hmm profiles from the multiple sequences alignments (hmmbuild 3.2.1 | June 2018 in HMMER 3.1b2 package February 2015) (http://hmmer.org/) that were subsequently aligned to our eukarya set using hmmsearch (version 3.2.1 | June 2018 in HMMER 3.1b2 package February 2015) with a cut-off e-value of 10^-3^.

### Measuring co-occurrence with phylogenetic profiles and (non) interacting proteins

For all orthologies we constructed phylogenetic profiles by defining the presence (1) and absence (0) of all orthologs in the 167 species for a given orthology. For evaluating (non-)interacting proteins we obtained multiple protein interaction datasets. For interacting proteins we used the human BioGRID interaction dataset that contains physical interactions between proteins [33] (version 3.5.172 May 2019). We filtered this set to obtain pairs found in at least five independent publications (PubMed ID’s) as a measure of how thoroughly these proteins were investigated and how amenable they are to high-throughput measurements.

We defined a pseudo negative interaction set from BioGRID by taking pairs of the proteins that were found to be interacting at least five times, but not with each other. We applied these criteria so that the negative set only contains proteins that were found in other interactions and thus exclude the possibility of the interaction not being observed due to a myriad of technical reasons. Additionally, we used a previously compiled negative interaction dataset [34] and cross referenced this set with interactions reported in BioGRID to remove recently found interacting proteins. Finally, we defined a random interaction set from pairs of random proteins. Each set was mapped back to the different orthologies using the corresponding human protein id.

To evaluate the (dis)similarities between phylogenetic profiles of interacting proteins we calculated the distances between profiles using 13 distance and correlation measures (Table 2). The correlation measures were converted to distances (1 - correlation). Distance measures that had values other than between 0 to 1 were converted by dividing by the maximum value found for that distance. We chose the cosine distance and the pseudo negative set which had the best or equally as good predictive power for further analysis (see Results and Supplementary Figure 2).

**Table 2.** Different distance (D) or correlation (C) measures used to calculate the distances between phylogenetic profiles of OGs.

| Distance or Correlation measure | Values interval | Conversion |
| --- | --- | --- |
| Braycurtis | [0,1] | - |
| Cityblock (Manhattan) | [0,∞] | $D / \max(D) = D$ |
| Cosine | [0,1] | - |
| Dice | [0,1] | - |
| Euclidean | [0,∞] | $D / \max(D) = D$ |
| Jaccard | [0,1] | - |
| Kendalltau | [-1,1] | $(1-C) / 2 = D$ |
| Kulsinski | [0,1] | - |
| Rogertanimoto | [0, 1] | - |
| Russellrao | [0,1] | - |
| Sokalmichener | [0,1] | - |
| Spearman | [-1,1] | $(1-C) / 2 = D$ |
| Yule | [0,∞] | $D / \max(D) = D$ |

Additional to the distance between the phylogenetic profiles of interacting proteins within a method, we calculated the cosine distances between phylogenetic profiles of orthologous groups of different orthology methods. The groups compared are the ones that mapped back to the same human gene, which ideally should be a similar/the same OG.

### Comparing against manually curated orthology sets and all-vs-all inference methods

To benchmark the automatically inferred OGs against a set of high quality OGs, we took manually curated sets of protein complexes constructed previously by members of our lab and mapped them to our latest dataset version. The manually curated orthology set contains a total of 125 OGs (Table 3).

**Table 3.** Manually curated OGs.

| Complex/Pathway | Number of OGs | Study/Reference |
| --- | --- | --- |
| Intraflagellar transport complex | 26 | (van Dam et al., 2013) |
| Kinetochore | 91 | (van Hooff et al., 2017; Tromer et al., 2019) |
| TBP-associated factors | 8 | (Antonova et al., 2019) |

We used two cluster overlap measures to compare the overlap between the manually curated and automatically inferred OGs. The the F-Grand K-Clique Score (FGKCS) [35] matches cliques of (in our case) OG members within the set of all possible cliques between OGs from automated methods and our manually defined set of OGs and determines the performance using the F-Grand metric.

The Adjusted Rand Score (ARS) [36], roughly, by counting how many pairs in each cluster occur together in the same cluster between methods and is adjusted for chance (and not on overlap/intersection like Jaccard similarity index). Because the ARS focusses on pairwise agreement between clusters, methods that define and/or generate singleton clusters (orthologous groups of size 1) are penalised more than methods that do not. To avoid over-penalising, we only compared LECA orthologous groups, thus removing singleton clusters from the analysis.

To better understand how ARS and FGKCS compare to actual cluster similarity between the inferred and manual OGs, we first stepwise shuffled a percentage of labels for one automated orthology definition and compared it with the same but unshuffled definition. We measured the relationship of ARS and FGKCS compared to the OG similarity as the percentage of unshuffled members.

To visualize how the manually curated OGs were represented in the automatically inferred OGs, we looked at the number of proteins overlapping between each manual OG and automatically inferred OGs. We divided this overlap number by the total amount of proteins in the manual OGs. This in turn gave us a matrix of overlap fractions that could be visualized for each manually vs. automatically inferred OG. We additionally calculated the percentage under- and oversplit sequences. We did this by counting the amount of sequences that were not in the OG containing the most sequences (under the assumption that this OG is the correct one) per row (oversplitting) or column (undersplitting). This number we divided by the total amount of sequences in this row or column. We calculated the recall of the manual OGs by taking the total amount of sequences assigned from the manual OGs to the automatically inferred LECA OGs and divided it with the total amount of sequences in the manual OG set (i.e. 5852).

## Results and Discussion

### Comparison between orthology methods show similar behaviour after inferring LECA orthologous groups

To investigate different orthology methods and how well they recapitulate a number of observations regarding genome evolution, we first inferred orthologies on a diverse set of 167 eukaryotic proteomes [18] using eight different inference methods or their settings. Supplementary Table 2 shows an overview of the statistics of every orthology inference method and their complete set of inferred orthologous groups (OGs).

In the original orthology definition, Fitch [37] described that if the history of a single gene is as the history of the species, they should be called orthologous, i.e. they are a single gene in the ancestor of the two species and resulted from a speciation there rather than a duplication earlier. Extrapolated for eukaryotic species this would imply that an inferred orthology relation for a set of diverse eukaryotes should equate to the presence of a gene in LECA. Despite labelling themselves as orthology methods, most methods might not explicitly check if the inferred OG indeed represents an ancestral presence. Hence, in addition to inferring the OGs with the methods as is, we estimated LECA OGs using a slightly more strict extension (see Methods and Materials) of the Dollo parsimony method [29–31] and calculated their (independent) loss to extant species.

Except for the outlier values of Ancestral Panther and SwiftOrtho, the inferred orthologies are similar between the methods while comparing the statistics for inferred LECA OGs (Figure 1). The high amount of automatically derived LECA OGs is consistent with previous large scale studies [30,32] and the observation from multiple manual analyses where individual OGs are found to be in fact in LECA OGs [38].

**Figure 1.**
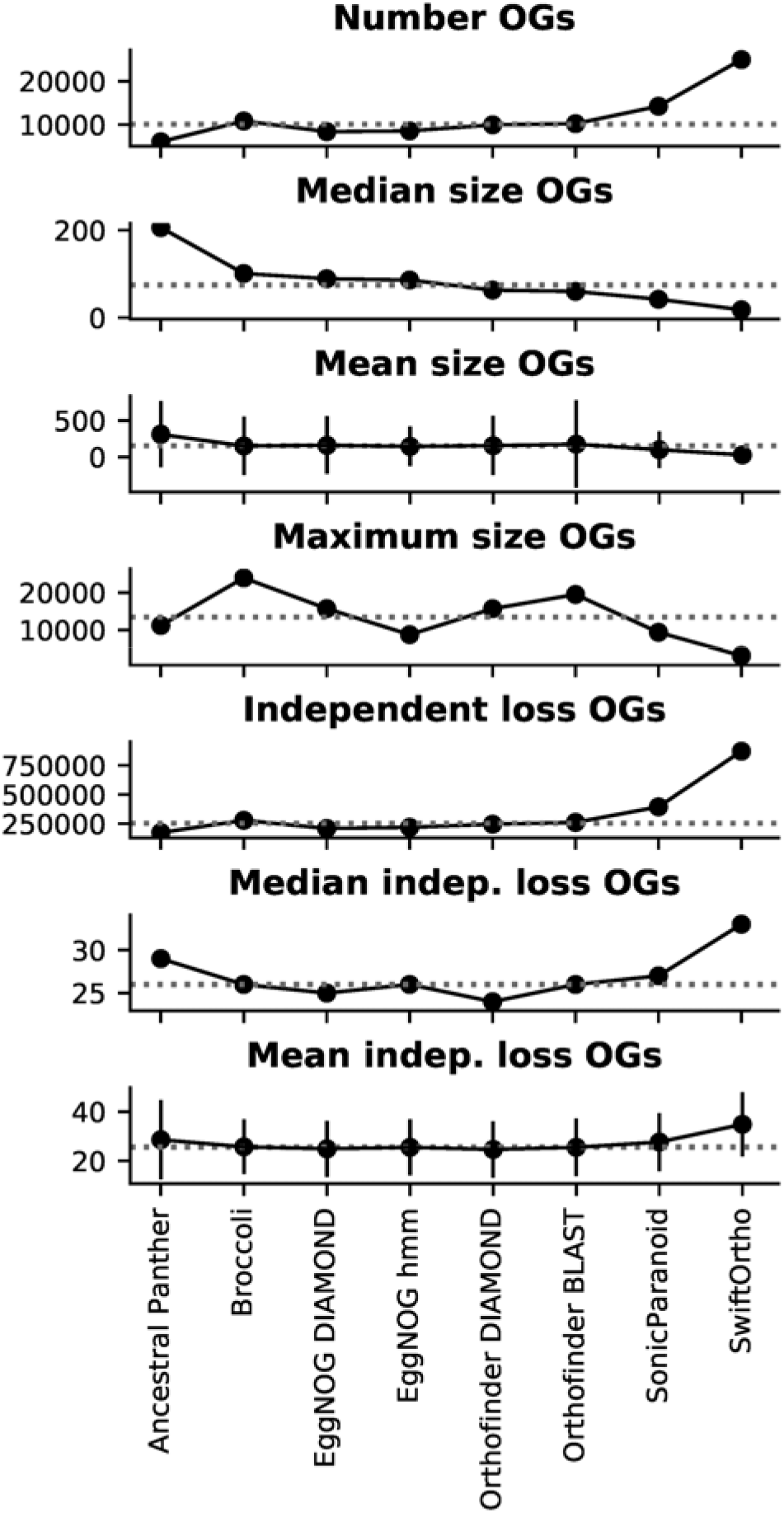
Comparison of different orthology methods after inferring LECA OGs. Shown are the statistics of the number of LECA OGs, size of the OGs and independent loss measures. Grey dotted line shows the median value of a given metric. The vertical lines in the mean size OGs and mean independent (indep.) loss OGs are standard deviations.

Independent loss distributions behave relatively similar between orthologies (Figure 2), with little differences in median, means and standard deviations (Figure 1), except for Ancestral Panther, SonicParanoid and SwiftOrtho. Ancestral Panther reports the lowest number of LECA OGs, but the median OG size is the highest of all the methods (Figure 1), indicating that the Panther OGs are very inclusive. This in turn results in a broader distribution of independent loss, and thus higher median loss, for these OGs. SwiftOrtho reports the highest number of LECA OGs, with a smallest median OG size (Figure 1), indicating SwiftOrtho is very strict. This inflates the independent loss distribution, giving SwiftOrtho the highest independent loss off all the methods.

**Figure 2.**
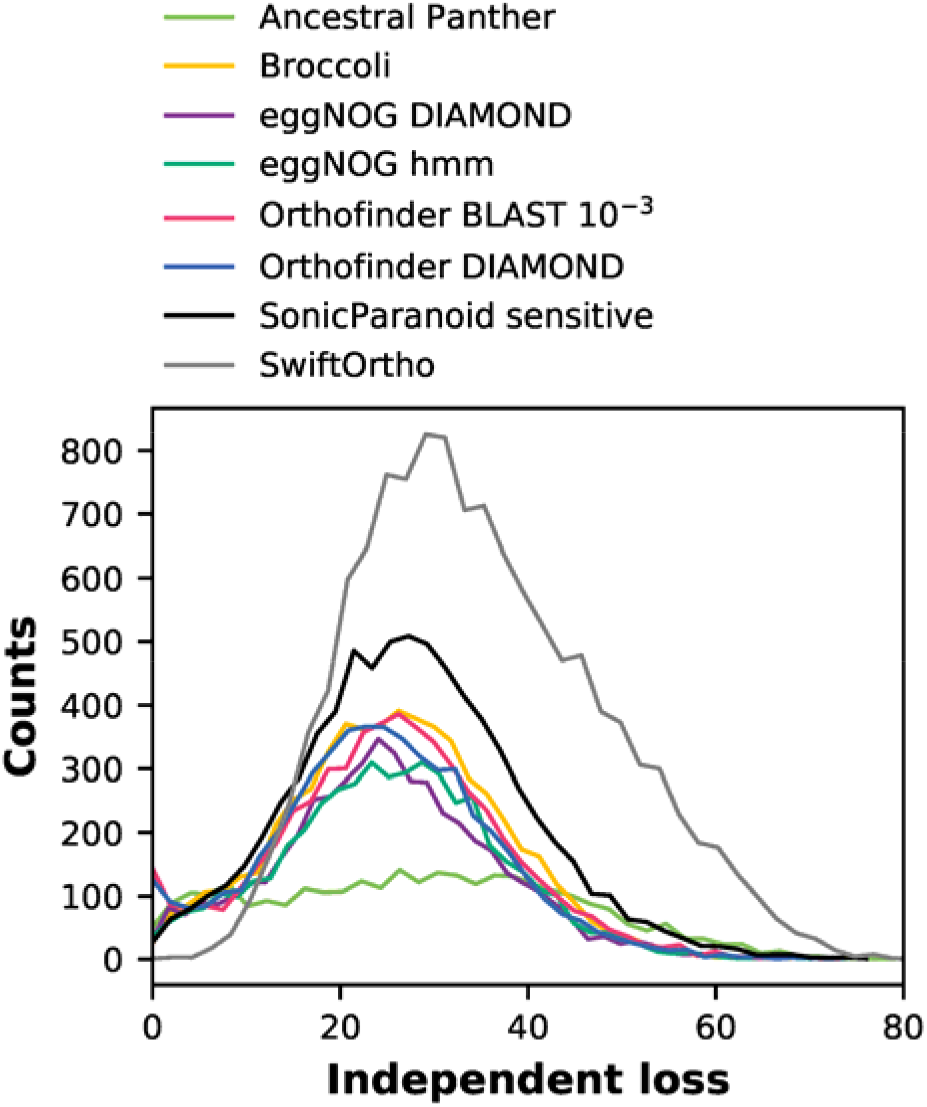
Loss distributions of LECA OGs estimated from different orthology inference methods. Except for Ancestral Panther, SonicParanoid and SwiftOrtho, the loss distributions show similar patterns. For eggNOG hmm vs. Orthofinder BLAST/Broccoli and eggNOG DIAMOND vs. Orthofinder DIAMOND, there is no significant difference between distributions (Kruskal-Wallis H test p-value > 0.001).

Nevertheless, the number of independent loss is high for all inferred orthologies, as is expected [30,31]. The number of independent loss has been shown to be influenced by gene prediction problems causing falsely inferred gene absences [18], where a suspicious absences can often be found back in the DNA. However, this is likely an equally big problem for all methods, since many orthology methods require predicted proteomes and cannot run on DNA/six frame translated DNA.

### Co-occurrence of interacting proteins is predicted similarly and fairly well by most of the orthology methods

Independent gene loss is not random [5]. In fact, there are countless observations that cooccurring (or co-lossed) proteins tend to interact [1–4,39]. This creates an additional opportunity for evaluating orthology methods, namely how well different methods can predict co-occurrence of interacting proteins. This metric is also of great relevance for phylogenetic profile methods [39]. For this reason, we calculated distances between phylogenetic profiles of LECA OGs for every method and assessed how well protein-protein interactions were predicted by that method.

These evaluations will identify orthology methods that are able to capture a high(er) degree of co-occurrence between interacting proteins and that are more capable of identifying ‘functional’ orthologous relationships over an arbitrary set of species.

We used the human BioGRID protein interaction dataset [33] as a positive interaction set, and constructed a negative interaction set from BioGRID by selecting proteins that are well studied (i.e. detected to be interacting at least with five other proteins), but have not been detected to interact with each other (zero interactions in BioGRID). We tested 13 distance measures (Table 2). For all distances we observed a clear signal for both the positive and negative interaction set (Supplementary Figure 1 and Supplementary Figure 2), with the exception of SwiftOrtho. The cosine distance had the best area under the curve (AUC) values compared to the other distances measured for most methods (Supplementary Figure 2).

There is no large difference in the predictive power for co-occurrence of interacting proteins from the different orthology inference methods (Figure 3). Orthofinder BLAST has a marginally higher AUC value than the other methods. SwiftOrtho has noticeable lower predictive power for co-occurrence.

**Figure 3.**
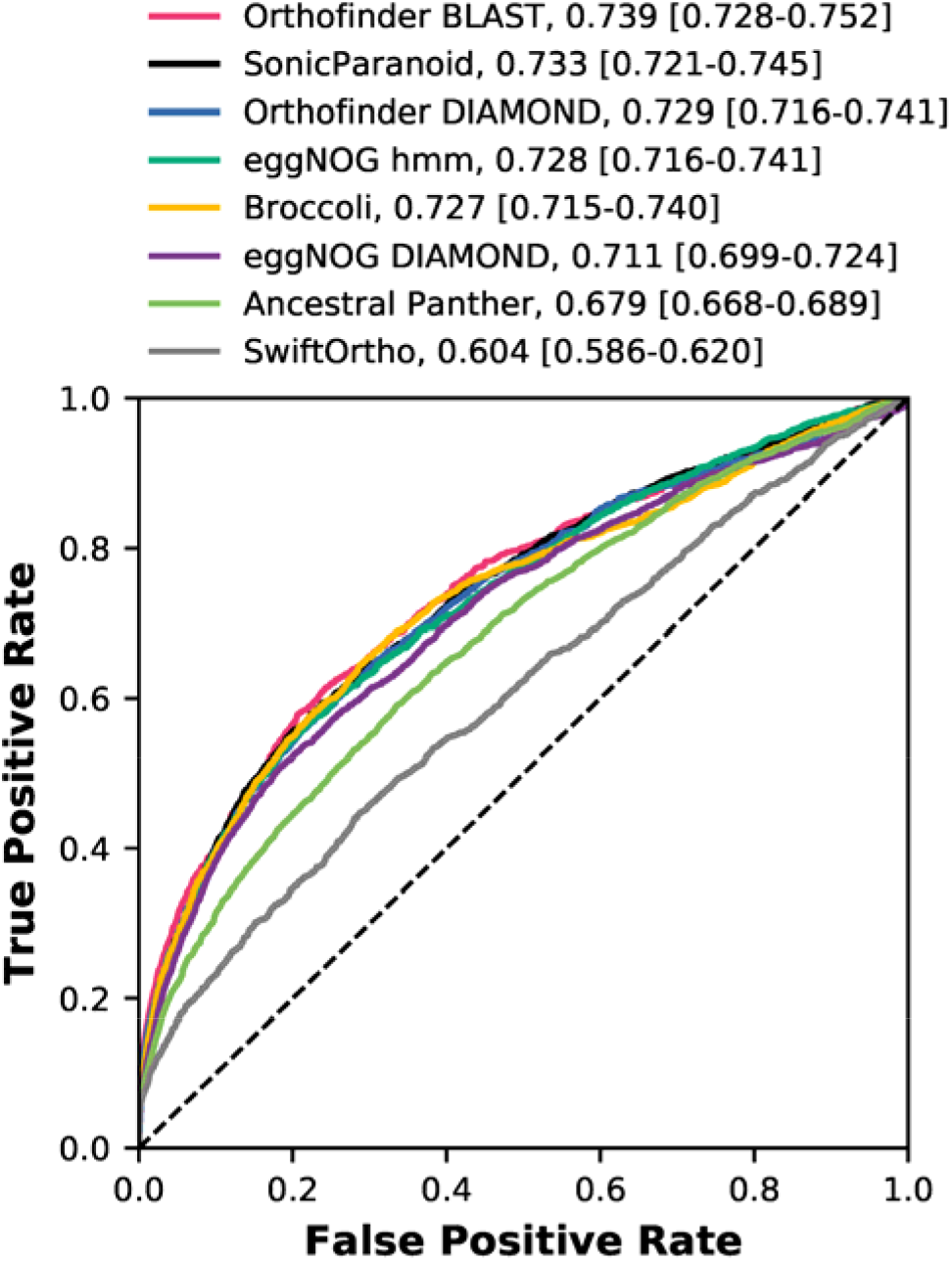
ROC plot comparing the predictive power of for co-occurrence of interacting proteins from different orthology inference methods, using the cosine distance. The legend additionally shows the AUC values for each curve and the confidence intervals (permutations n = 1000).

The quality of our predictions (as measured by AUC) is higher compared to a more elaborate phylogenetic profile method dedicated paper [40], combining phylogenetic profiles in conjunction with the MinHash technique (AUC value of 0.6155 for their human protein interaction set). Most importantly, most orthology methods successfully recapitulate the observation that loss is not random but co-occurs between interacting proteins.

A logical (and desirable) explanation for the similar performance between methods would be that human proteins are assigned similar phylogenetic profiles across different methods. However, when comparing OGs mapped to the same human protein. the phylogenetic profiles generated from the OGs by the different methods differed substantially from each other (Supplementary Figure 3.).

### OGs inferred by different methods show imperfect overlap in between methods and manually curated OGs

From previous work [1–4] we collected a set of 125 manually curated OGs at LECA level (Table 3). The comparison to manually curated OGs aligns our evaluation with other evaluation strategies, such as Quest For Orthologs, while the comparison of the inferred OGs to one another gives us a view of how similar the OGs are to each other.

The comparison of the inferred OGs with the manually curated OGs shows overall an imperfect, but decent, overlap for all methods (Figure 4 for Adjusted Rand Score (ARS), and Supplementary Figure 4 for F-Grand K-clique Score (FGKCS)). The overlap between the manual set against all inferred OGs is very high (mean ARS of 0.85) compared to the all-vs-all comparisons (mean ARS of 0.52) between the inferred OGs. This is due to the fact that the manual OG set is a smaller subset of sequences and thus more easily included wholly into the inferred larger OGs, but also because errors in the inferred OGs are measured one-sided to the manual OGs, compared to the all-vs-all comparison where errors are measured two-sided. To understand how the ARS relates to actual OG similarity between the manual and automatically inferred OGs see Supplementary Figure 5. (Supplementary Figure 6 for FGKCS).

**Figure 4.**
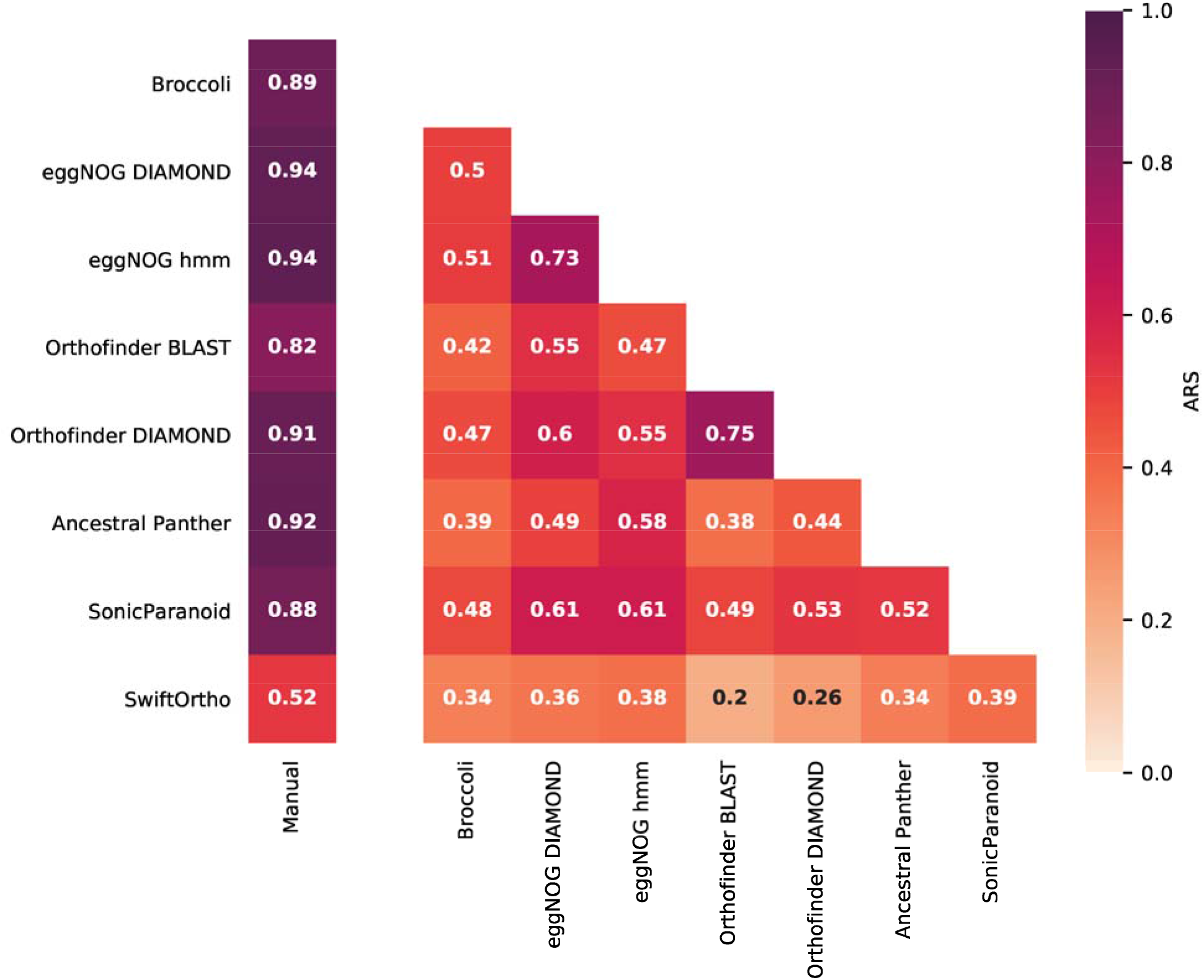
All-against-all comparisons of the different orthology methods and the manual set using the (Adjusted Rand Score) ARS. An ARS of zero would indicate that two OGs vary at a level that would be expected by chance, while an ARS of 1 is a perfect OG overlap.

An important and rather problematic observation is that the comparison between the inferred OGs of different methods shows there is low overlap, or little consistency, between the methods. As expected, similar methods have similar OGs (eggNOG hmm vs. eggNOG DIAMOND and Orthofinder BLAST with Orthofinder DIAMOND). Nevertheless, even between these methods there is still a considerable difference. It is important to uncover which types of errors are made in each automated method.

### Representation of the manually curated OGs in automatically inferred OGs - where are the differences?

In order to investigate what types of errors are made by the different orthology methods, we wanted to see how the various manually curated OGs are represented in the inferred OGs. We visualized the fraction of proteins per manual OG overlapping with matching proteins assigned to the inferred OGs of the different methods, to the total amount of manual proteins in a given manual OG (Figure 5). Multiple grid points in the same column mean that multiple manually curated OGs are contained within a single automatically inferred OG, indicating an undersplitting in the orthology inference, due to for instance unrecognized (pre-LECA) (out-)paralogs. A clear example of undersplitting is Cdc20 and its outparalog Cdh1 arising from a, by most of the the methods unrecognized, pre-LECA duplication [4] that as a consequence are consistently lumped together in the automated orthology inference methods in various degrees (Figure 5 and Supplementary Figure 7). The same applies for the mitotic kinases Aurora and Plk.

**Figure 5.**
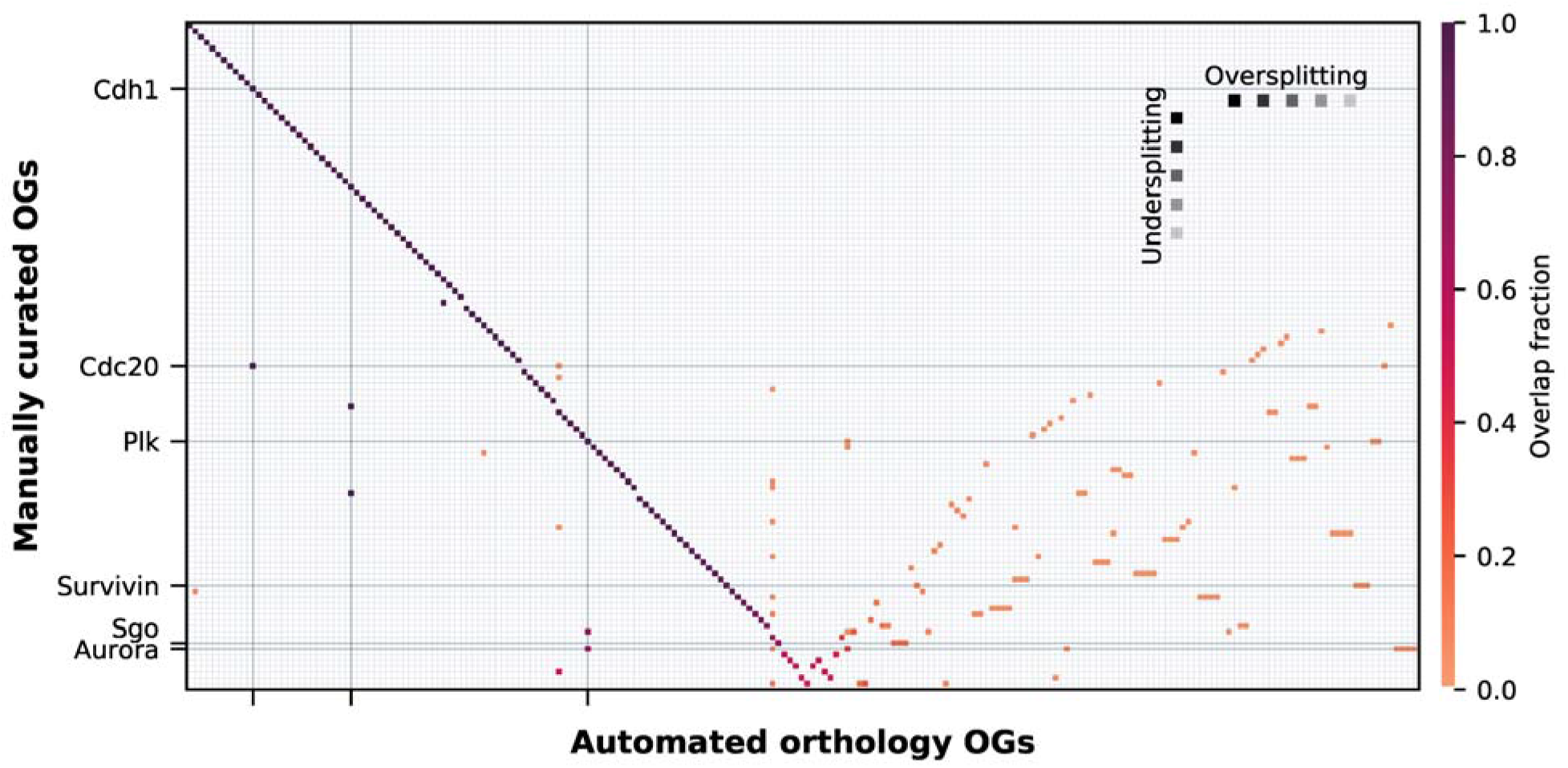
Heatmap showing the fraction of overlap of the clusters between the manually curated OGs (only examples are labelled) and the inferred OGs from, in this case, Orthofinder BLAST as example. The manually curated OGs (y-axis) are sorted from highest to lowest overlap with an Orthofinder OG (x-axis), creating a diagonal (dark purple). The non-diagonal part (lower right corner) is clustered with the weighted clustering method. On the left side of the diagonal there are clear examples of undersplitting of the manually curated OGs. To the right of the diagonal (lower right corner) there are clear examples of oversplitting, or misclassifications. The colorbar shows the fraction of overlap between the manual OG sequences to the automated OG sequences.

Multiple grid points in the same row of Figure 5 means that a single manually curated OG is contained within multiple inferred OGs, indicating an oversplitting in the automated orthology, due to for instance misclassification, unrecognized homology, or lineage specific duplications (in-paralogs). A few examples of oversplitting are of the OGs Aurora, Cdc20, Plk, Sgo, Survivin (Figure 5) and MadBub (Supplementary Figure 7), which are consistently fragmented to a higher degree in most automated orthologies (Supplementary Table 3). These proteins have elevated copy numbers, due to possibly recurrent duplications (and subfunctionalization) [2].

The bulk of the automated OGs assigned correctly to the manually curated OGs (diagonal with dark purple grid points (Figure 5). Although, under- and oversplitting can be seen in different degrees between the methods. To compare the degree of under- and oversplitting of the manual OGs between automatically inferred ortholgies we calculated the percentage of sequences that are under- and oversplit in the different methods (Figure 6). This shows that, overall, SwiftOrtho has the least undersplitting, but has the lowest recall and most oversplitting of all the methods. EggNOG hmm and eggNOG DIAMOND have the least undersplitting and Orthofinder DIAMOND the least oversplitting. These values indicate that eggNOG is good at recognizing more distant homology, while Orthofinder DIAMOND is better in correctly classifying orthologs, and detecting recent duplication, but the numbers between the methods are fairly similar. Looking at the total amount of manual OG sequences that are assigned to a LECA OG in the automatically inferred orthologies, we see that Ancestral Panther has the highest percentage of manual OG sequence recall.

**Figure 6.**
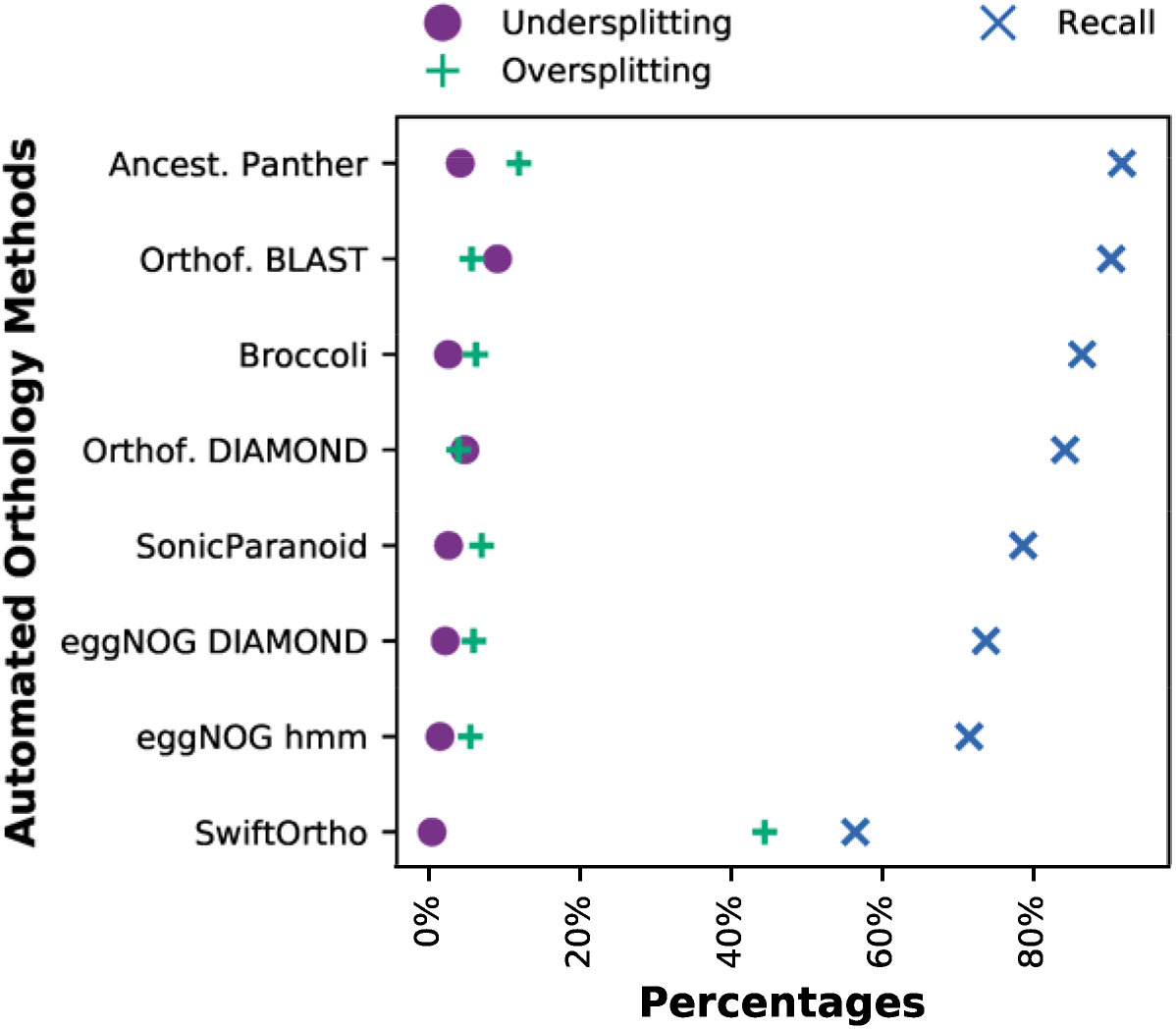
Degree of under- and oversplitting of the manual OGs in the different orthology methods. The methods are sorted from highest (top) to lowest (bottom) recall of the manual OG sequences in the automated orthologies.

## Conclusions and outlook

Orthology prediction is difficult, but pivotal, in comparative genomics. Many methods and tools have been developed to capture orthologous relationships between genes as accurately as possible. We evaluated several orthologies created by automated orthology methods with different underlying algorithms for their ability to capture a number of key observations on eukaryotic genome evolution.

We show that co-occurrence of interacting proteins is predicted similarly well by all orthology methods that were tested and most are similar in behaviour when determining gene loss patterns in eukaryotic evolution. However, they show imperfect overlap with manually curated OGs. Important to note is that, although all orthology inference methods used in this study are similar when describing general patterns in eukaryotic genome evolution, they show large differences amongst the inferred orthologies themselves.

Different methods can provide more optimal solutions compared to others when studying different evolutionary scenarios, such as coevolution, grouping more distant orthologs or more recent duplications. However, many challenges remain in orthology inference, both biological (orthology vs. paralogy) and practical (data increase and computational resources), all of which we have experienced throughout this study. Nevertheless, our results show that (automated) orthology methods show similar behaviour with respect to large scale evolutionary observations, such as loss patterns, but caution is warranted when looking at the smaller scale, such as single OGs of interest.

Automatic and manual orthology methods are complementary. Leveraging this complementarity could improve comparative genomics in the near future. This means that (orthology) databases should aim to provide OG assignments in a convenient way to accommodate manual analyses and vice versa to be able to use manual assigned OGs as seeds in automated orthology analyses, or to guide development of orthology inference tools.

## Funding

This work was supported by the Netherlands Organisation for Scientific Research as part of the research programme Innovational Research Incentives [016.160.638].

## Acknowledgements

We would like to thank the developers of Ancestral Genomes, Xiaosong Huang and Paul Denis Thomas, for providing us with the additional data used in this study. We would also like to show our gratitude to Salvatore Cosentino, who provided us with the SonicParanoid results on our dataset.

## Notes

https://github.com/ESDeutekom/ComparingOrthologies

### Competing Interest Statement

The authors have declared no competing interest.

