## Supplementary Figures for "Comparing orthology methods and their performance by recapitulating patterns of eukaryotic genome evolution"

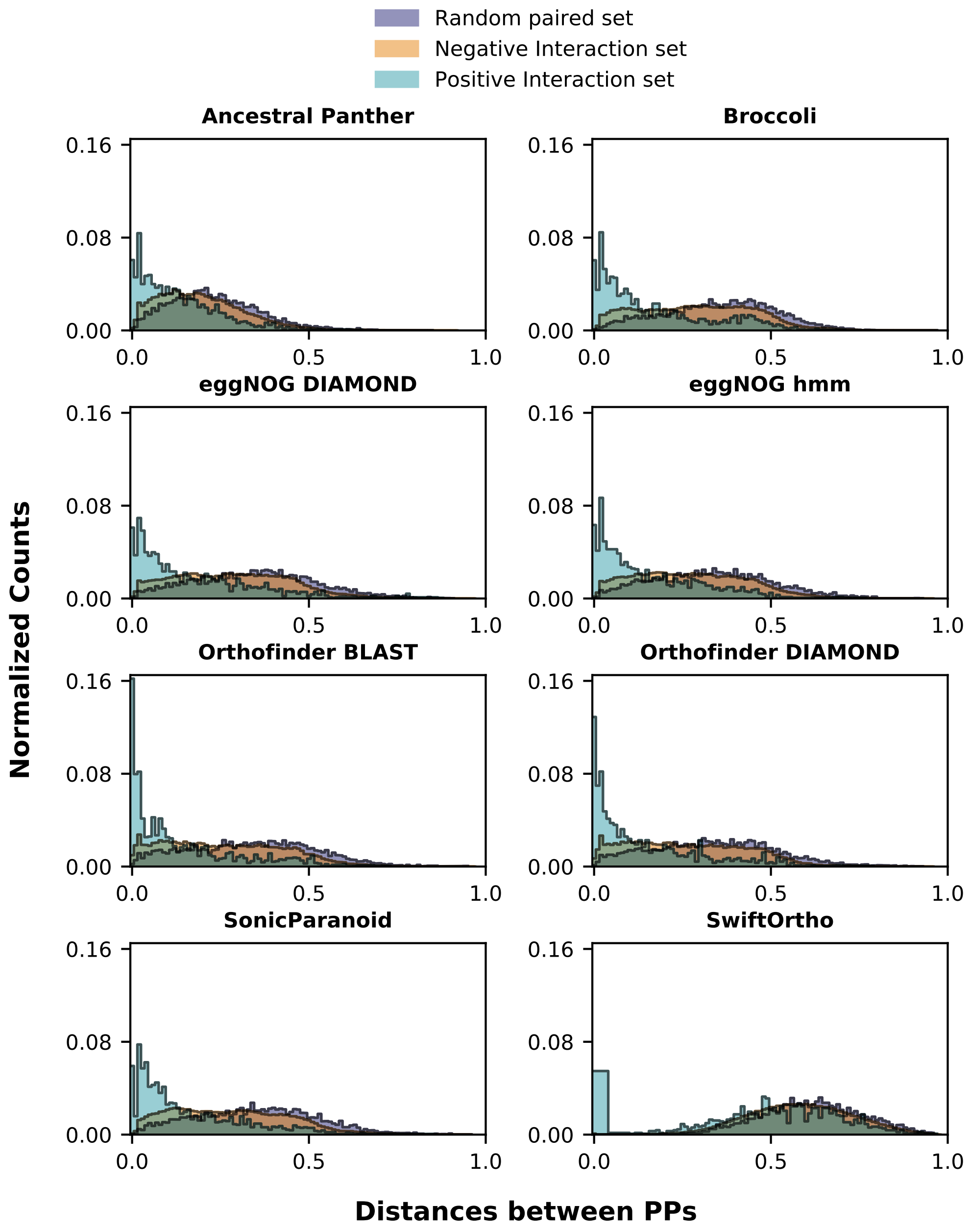


**Supplementary Figure 1. Distribution of cosine distances between phylogenetic profiles.**

Example distributions of the cosine distances between phylogenetic profiles of the positive (blue), negative (orange) and random (purple) protein interaction sets, showing there is a
(different) signal in de phylogenetic profile distances for the positive and negative interaction set, compared to the distances of a random protein interaction, for all but SwiftOrtho. The counts are normalized by the sum of all counts.

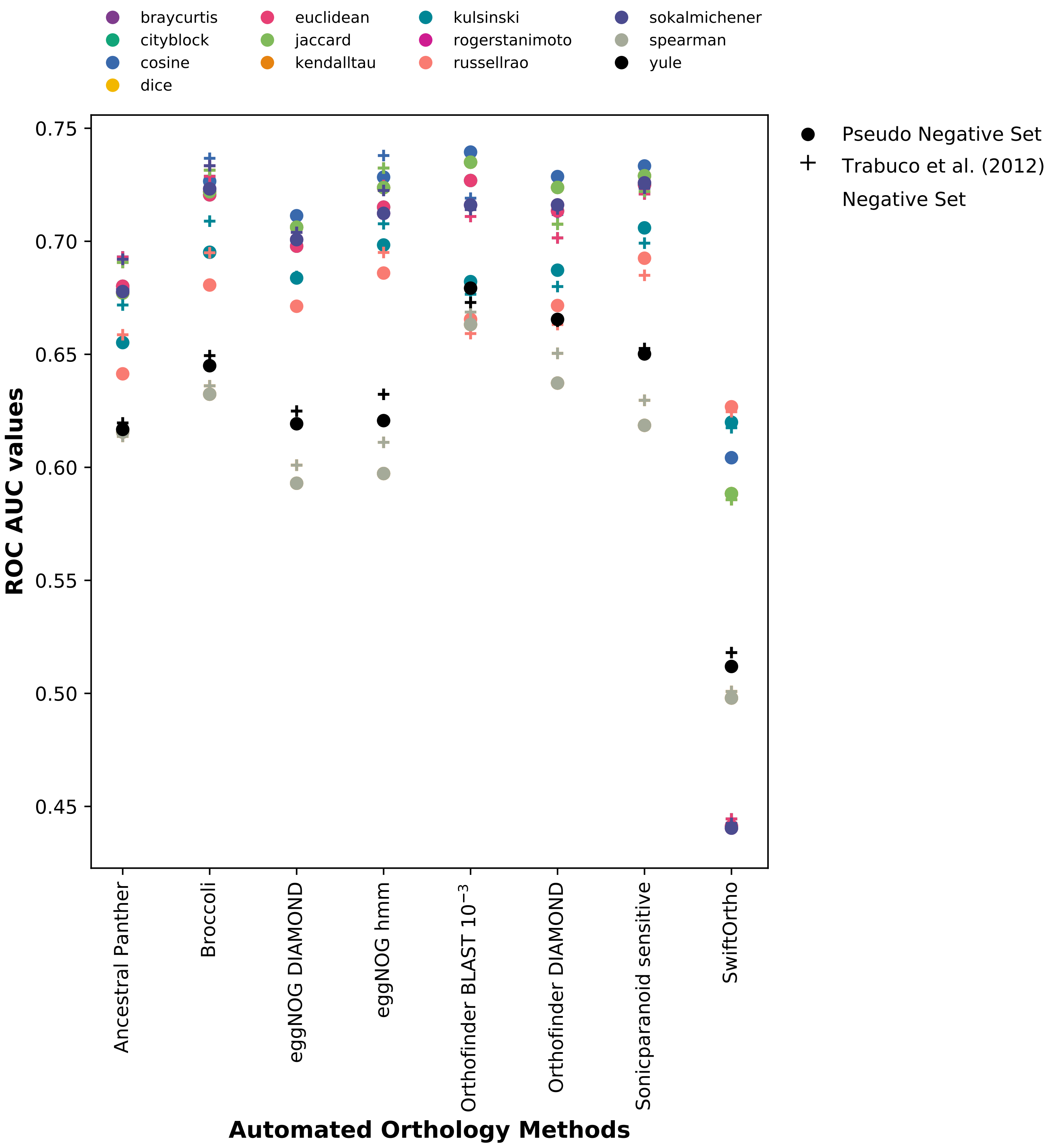


**Supplementary Figure 2. Comparison between multiple methods using multiple distances and interaction sets.**
Calculated ROC Area Under the Curve (AUC) values are shown for all the different methods. Cosine gives the highest AUC value for, except Ancestral Panther and SwiftOrtho, all methods. The Pseudo negative interaction set outperforms the Trabuco et al. (2012) negative interaction set 5 out of 8 times.

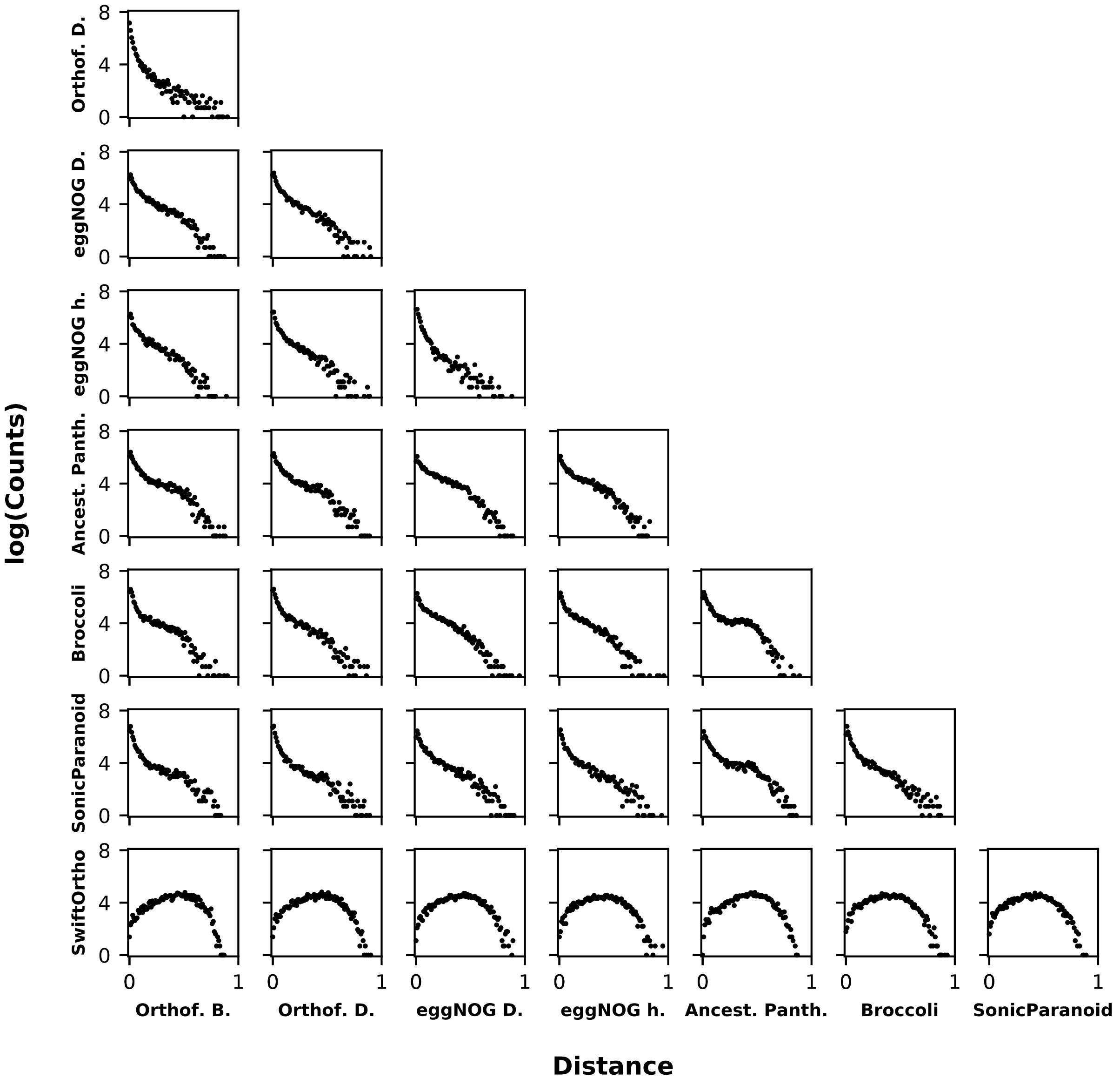


**Supplementary Figure 3. The cosine distances between the phylogenetic profiles of OGs from different orthology inference methods.**

The distances between the phylogenetic profiles of the orthologous groups mapped between the methods should ideally be similar, i.e. a human protein should be found in similar OGs between the different methods. However, this figure shows diversity between the phylogenetic profiles and thus the inferred OGs.

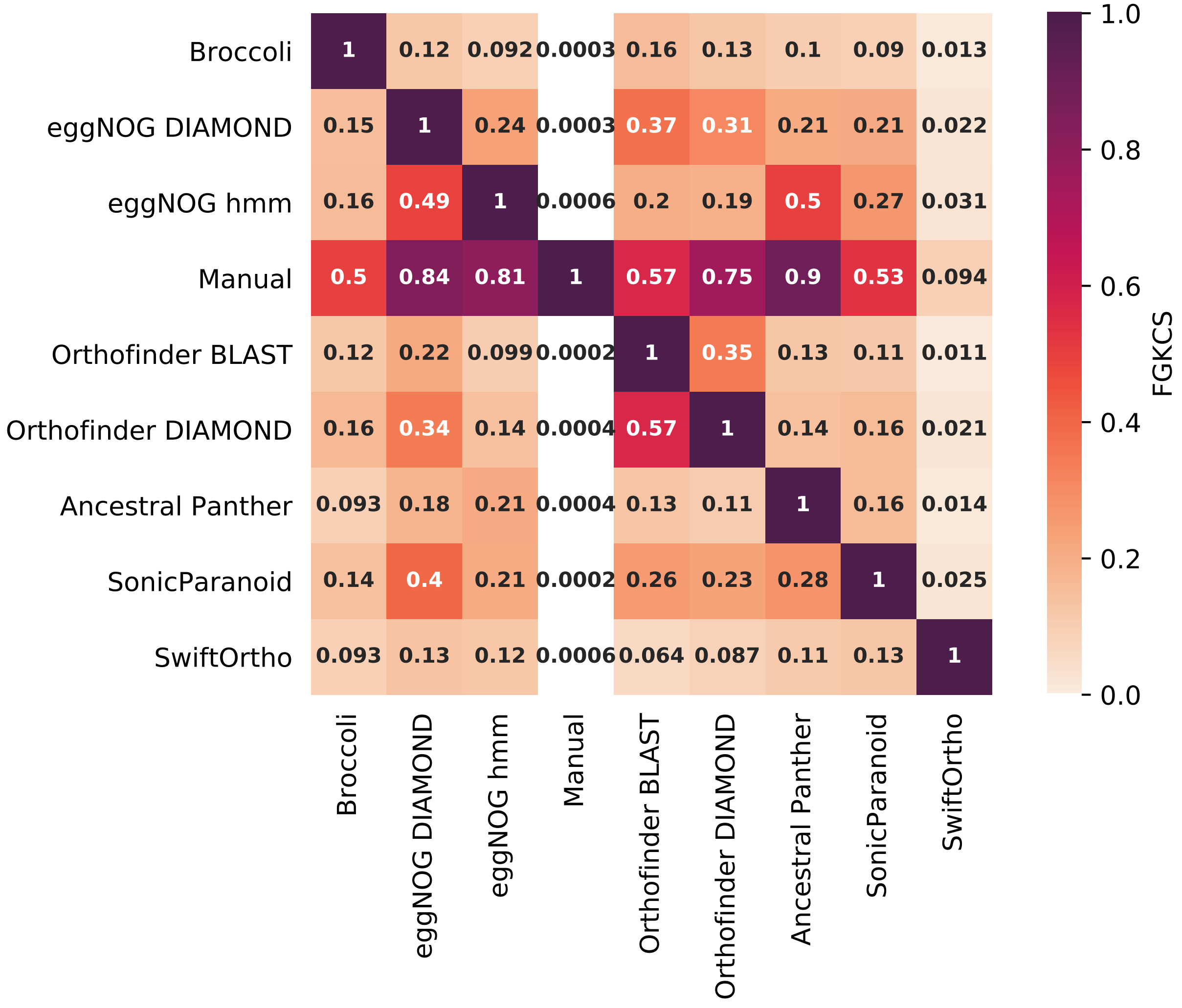


**Supplementary Figure 4. The F-Grand K-clique Score (FGKCS) score heatmap.** Unlike the ARS (main Figure 4), the FGKCS is not a symmetric score. This way we see how the OGs overlap from method A (vertical axis) to B (horizontal axis), and B to A. For instance, we can make out that the manual OGs (on vertical axis) clusters better with the inferred OGs (on horizontal axis), than the inferred OGs (on vertical axis) cluster to the manual set (on horizontal axis). This has to do with the OG size imbalance between the manual OGs and the inferred OGs (see main text Results).

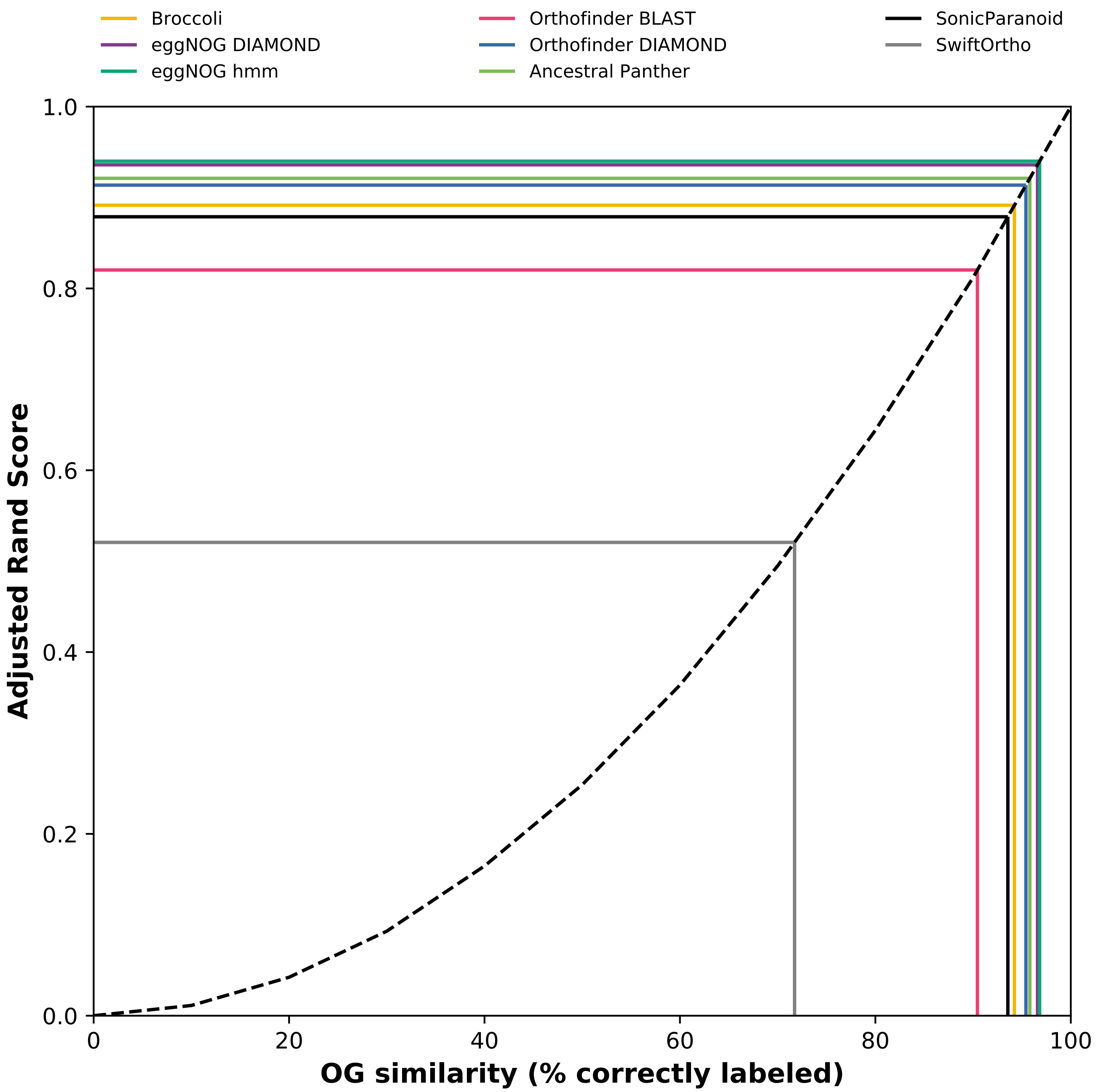


**Supplementary Figure 5. The Adjusted Rand Score (ARS) compared to OG similarity.** Shows (black dashed line) the relationship of ARS compared to the OG similarity measured as the percentage of unshuffled members (see Methods and Materials main text). The general shape of this relationship is non-linear. The ARS for all the orthology definitions are plotted (colored lines) to approximate the percentage of correctly assigned labels from the manually curated OGs by the automated orthology. The ARS shows a clear difference between the low cluster overlap of SwiftOrtho and the (lower) overlap of Orthofinder BLAST.

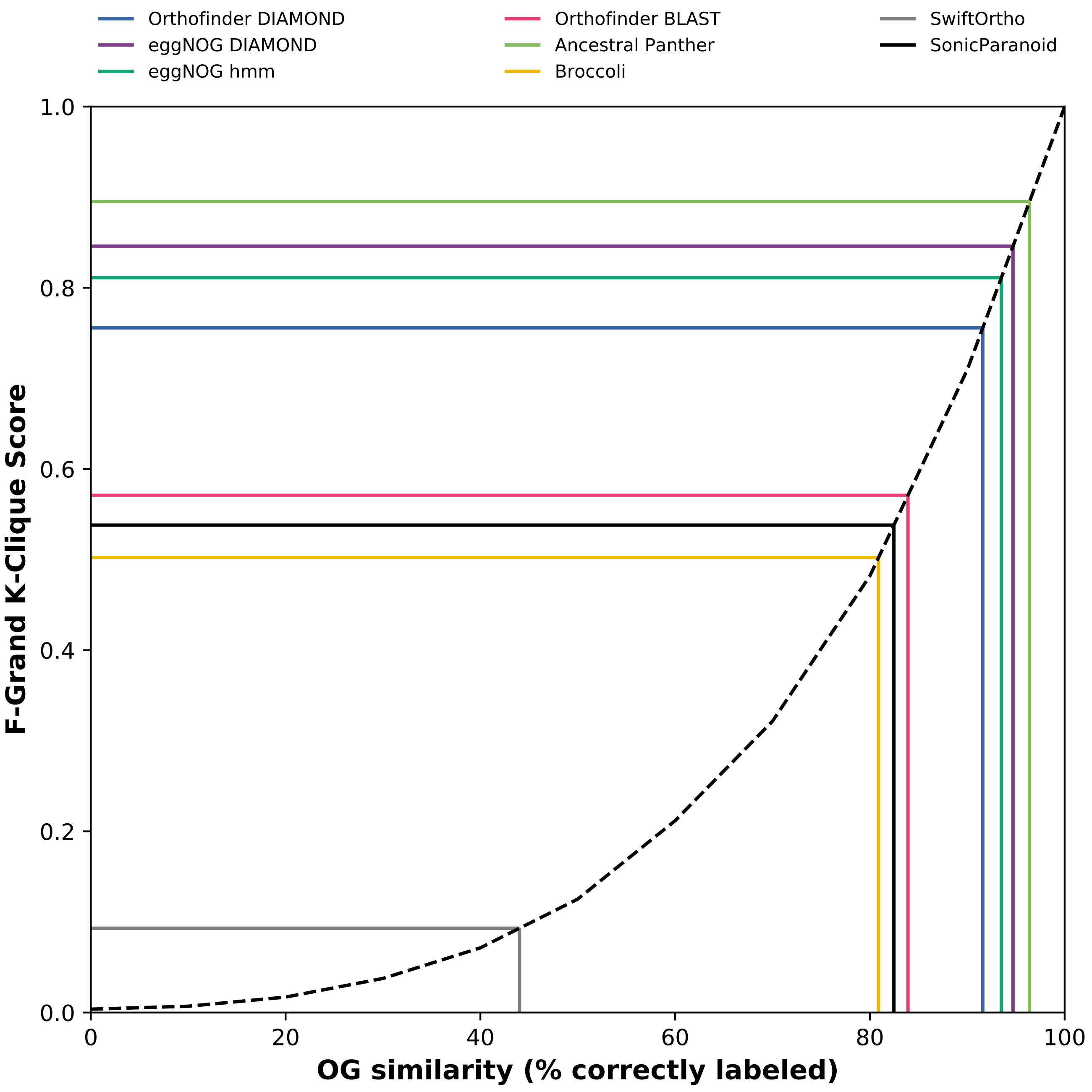


**Supplementary Figure 6. The FGKCS compared to OG similarity.**
Shows (black dashed line) the relationship of FGKCS compared to the OG similarity measured as the percentage of unshuffled members (see Methods and Materials main text). The general shape of this relationship is non-linear. The FGKCS for all the orthology definitions are plotted (colored lines) to approximate the percentage of correctly assigned labels from the manually curated OGs by the automated orthology. The FGKCS shows a clear difference between the (lower) cluster overlap of SwiftOrtho, second tier overlap of Broccoli, SonicParanoid and Orthofinder BLAST and the best cluster overlap for the rest of the orthologies.

**Supplementary Figure 7: The fraction of overlap of the clusters between the manually curated OGs and the inferred OGs from different orthology inference methods.**

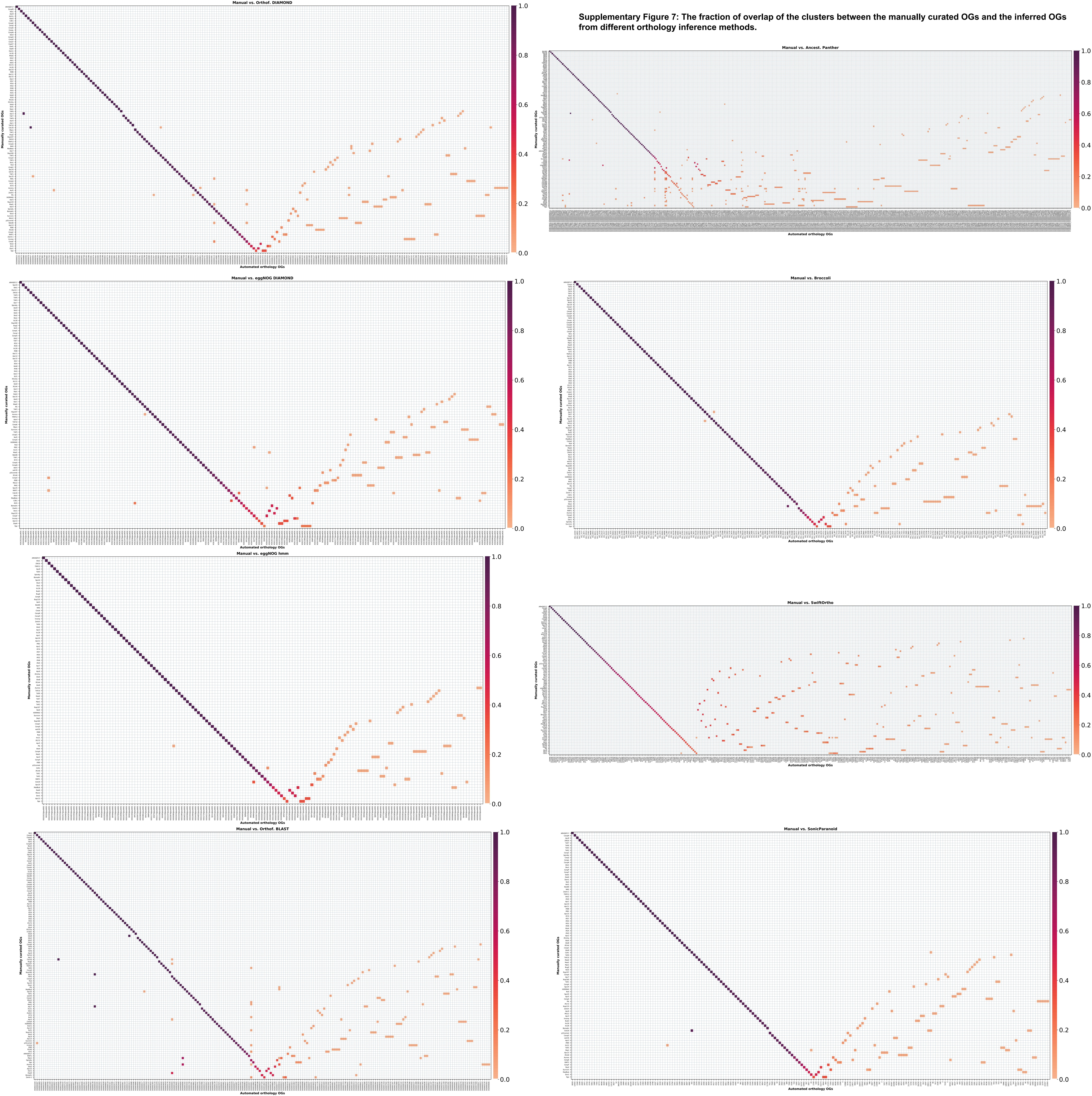
